# Emergent low-frequency activity in cortico-cerebellar networks with motor skill learning

**DOI:** 10.1101/2022.05.18.491978

**Authors:** Pierson J. Fleischer, Aamir Abbasi, Andrew W. Fealy, Nathan P. Danielsen, Ramneet Sandhu, Philip R. Raj, Tanuj Gulati

## Abstract

The motor cortex controls skilled arm movement by recruiting a variety of targets in the nervous system, and it is important to understand the emergent activity in these regions as refinement of a motor skill occurs. One fundamental projection of the motor cortex is to the cerebellum. However, the emergent activity in the motor cortex and the cerebellum that appears as a dexterous motor skill is consolidated is incompletely understood. Here, we report on low-frequency oscillatory (LFO) activity that emerges in cortico-cerebellar networks with learning the reach-to-grasp motor skill. We chronically recorded the motor and the cerebellar cortices in rats which revealed the emergence of coordinated movement-related activity in the local-field potentials (LFPs) as the reaching skill consolidated. We found that the local and cross-area spiking activity was coordinated with LFOs. Finally, we also found that these neural dynamics were more prominently expressed during accurate behavior. This work furthers our understanding on emergent dynamics in the cortico-cerebellar loop that underlie learning and execution of precise skilled movement.

## Introduction

The primary motor cortex (M1) is viewed as a driver for movement and an emerging view posits transient oscillatory dynamics-both at the level of spiking and local field potentials (LFPs) as the neural substrate for it^1–7^. There has been a particular interest in low-frequency quasi-oscillatory activity (LFOs) in M1, which can be brief (1-2 cycles) for rapid movements or longer for sustained movements, and it has been shown to be phase-locked to sub-movement timing^2,3,8,9^. Recent work showed that such oscillatory dynamics are coordinated in the M1 and dorsolateral striatum in the rodents as they learned a reach-to-grasp task^3^. One of M1’s principal projections is to the cerebellum via the pons^10–13^, but similar oscillatory dynamics have not been studied in cortico-cerebellar networks.

M1 is a key brain hub involved in voluntary forelimb movement: experimental lesions of M1 in animal models or neurological injury to M1 (such as stroke) impair dexterity^2,14–16^, stimulation of M1 neurons evokes movement^17–19^, spiking activity in M1 is closely linked to movement parameters^3,11,20–23^ and optogenetic perturbation of M1 affects forelimb behaviors^18,20,24,25^. The cerebellum’s role in the coordination of arm movements has also been extensively studied. Investigation of prehension/reaching tasks in animals have shown that cerebellar neurons – both in the cerebellar cortex and its deep nuclei are tuned to several movement-related events such as movement onset, cues leading to movement and its duration, limb position, velocity and muscle activity^26–31^. Besides coding for the above-listed features of limbs and associated movement parameters, other evidence indicates that the cerebellum participates in the formation of procedural memories, learning and retention of skills, habits, and conditioned responses^32,33^. Cerebellar lesions impair acquisition of skilled behaviors and patients with cerebellar disease show impaired reaching^34–36^. Furthermore, optogenetic perturbation of cerebellar nuclei or pontine inputs can cause a loss of endpoint precision in mice during reach-to-grasp behavior^13,37^. Additionally, electric stimulation over the cerebellum facilitates adaptive control of reaching^38,39^. Recent rodent work using two-photon imaging showed the emergence of shared neuronal-dynamics in *M1*-cerebellar ensembles as animals learned to expertly control a manipulandum^11^.

In this study, we have focused on transient oscillatory dynamics that emerge in M1 and the cerebellum as a reaching skill is learned. We recorded neural activity in the M1 and contralateral cerebellum (the primary M1 target through pons nuclei) throughout the learning of a reach-to-grasp skill in rats. We observed emergent coordinated low-frequency oscillatory activity (1-4 Hz) across M1 and cerebellum LFPs that was linked to increased success rates. We also found that LFPs modulated spiking in both regions and that the spiking dynamics were conserved for successful, accurate movements.

## Results

We trained 13 rats on the Whishaw forelimb reach-to-grasp task^40,41^ in our in-house built automated training box that is compatible with electrophysiology (**Fig. 1A**)^2,41,42^. We chose this task due to its similarity to skilled learning tasks in humans^43,44^ as well as extensive evidence that this task is associated with multiple levels of neural plasticity in the M1 and the cerebellum. Examples of this include changes in Long-Term Potentiation (LTP)^45^, dendritic spine growth^46^ motor map plasticity in the M1^47^, as well as patterned spiking in the cerebellar cortex^29^ and more recently, it has also been demonstrated that cerebellar associative learning underlies reach adaptation^48^. Importantly, patients with neurologic injury in either region show impairment in this skilled reaching behavior^36,49^. In a subset of rats (*n* = 5) that were monitored during reach-to-grasp motor skill consolidation, we also recorded neural signals, including single-unit activity and local field potentials (LFPs) in M1 and cerebellum (**Fig. 2A**). For the electrophysiology experiments, microelectrodes were implanted (microwire arrays in M1 and tetrodes/ polytrodes in cerebellum, see Materials and Methods; and **Supplementary Table 1**). In the animals that were recorded, training began five days after electrode placement surgery.

**Figure 1.**
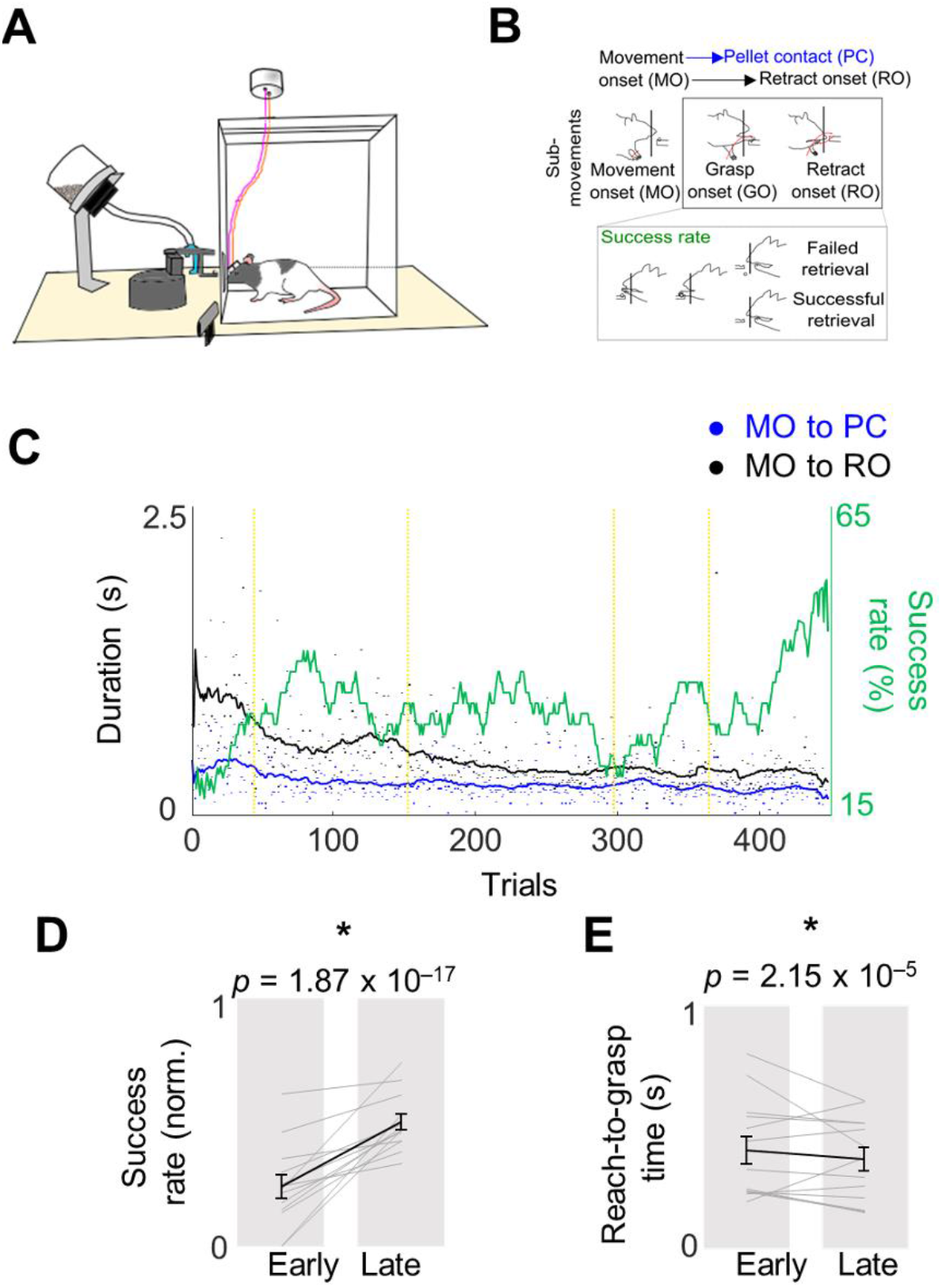
Behavioral evaluation of the skilled reach -to-grasp task.***A***, The behavioral setup for the skilled forelimb reaching task with simultaneous neurophysiological recording. ***B***, Top: Illustration of the reach-to-grasp task showing the three major parts of the reach movement: reach onset, pellet contact, and retract onset. Bottom: Comparison of a failed trial and a successful trial. ***C***, Success rate and reach event timing from a sample animal. ***D***,***E***, Difference in success rate and reach duration from early training days to late training days (n = 13 animals). Gray lines represent individual animals, and the black line is mean and s.e.m. across animals. P-values are from mixed-effects models.

### Measurement of Skilled Motor Performance

As in other studies that employ the Whishaw forelimb reach-to-grasp task, we assessed motor skill learning across two dimensions: speed and accuracy (**Fig. 1B-E**)^3,50^. Accuracy was measured as percent success in retrieving the pellet and speed was assessed using the time the animal took to perform the full reach-grasp-retract motor sequence. Training lasted for 5 days in automated behavioral boxes^2,41^, and animals performed 100–160 trials each day. Consistent with past results^3,50^ over 5 days of learning, the success rate increased and movements became faster as measured through reach-grasp task completion duration or reach duration (see **Fig. 1**). On average, success rates increased from 23.9 ± 4.7% to 49.8 ± 3.2% from early to late days (mean ± s.e.m., mixed-effects model: *P* = 1.87 × 10^−17^) and reach duration came down from : 406 ± 57 ms on early days to 367 ± 48 ms on late days (mean ± s.e.m., mixed-effects model: *P* = 2.15 × 10^−5^).

### Coordinated movement-related activity emerges across M1 and cerebellum during skill learning

We next evaluated the cerebellum in search of transient low-frequency oscillatory (LFO) dynamics similar to those that were recently shown to emerge in the M1^2,3^ while learning this skill. We observed that coordinated LFO (1–4 Hz) activity appeared during movement across M1 and cerebellum as performance improved (**Fig. 2**). This low-frequency activity was clearly observed in movement-related LFP signals (**Fig 2B, C**). The movement-related LFP power in the LFO-band increased from early to late days in both M1 and cerebellum (**Fig. 2C**; M1 baseline-normalized power: 0.51 ± 0.15 on early days to 0.65 ± 0.16 on late days, mixed-effects model: *t*(406) = 4.3, *P* = 2.3 × 10^−5^; cerebellum power: 0.43 ± 0.12 to 0.86 ± 0.25, mixed-effects model: *t*(650) = 8.6, *P* = 8.4 × 10^−17^).

**Figure 2.**
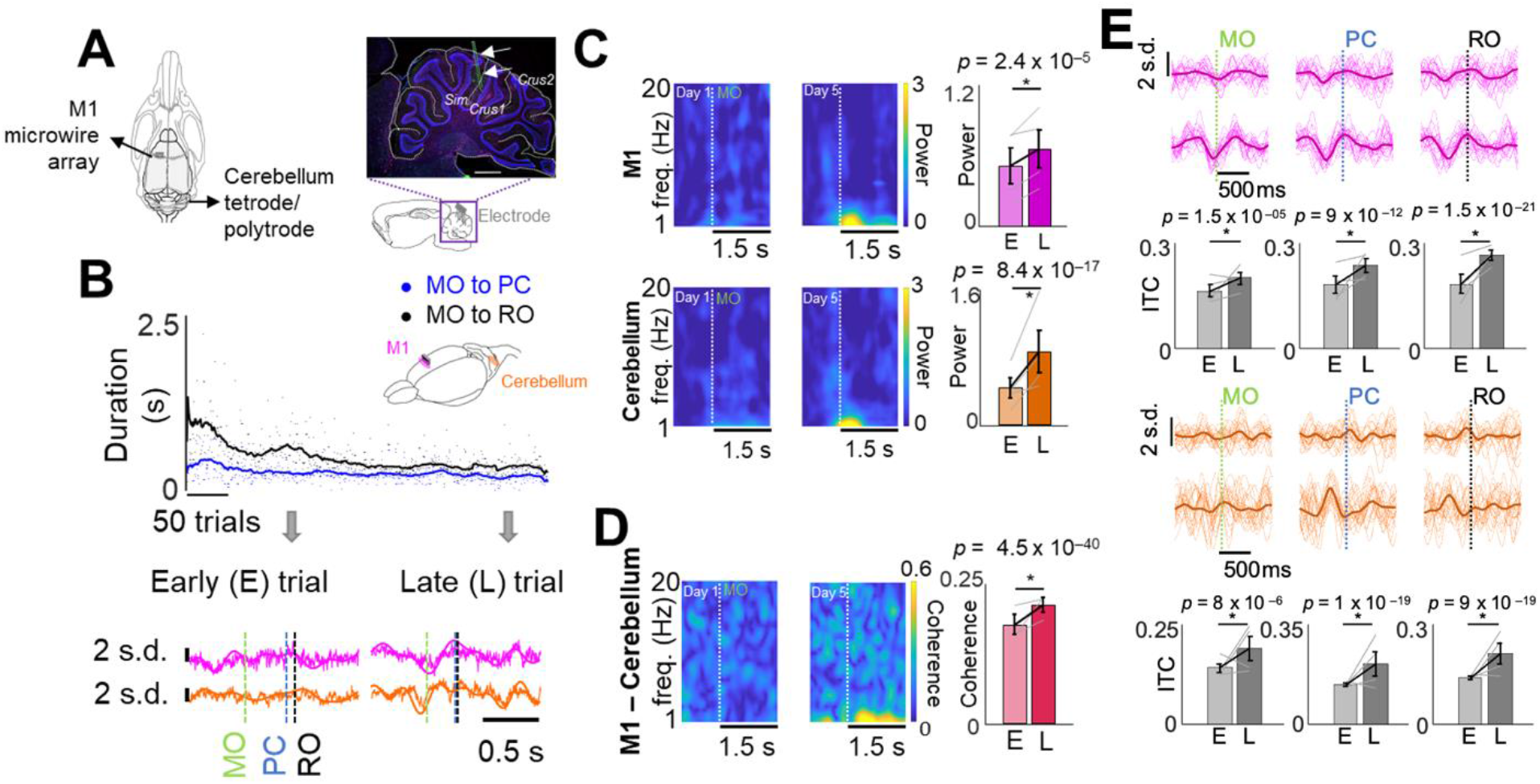
Coordinated movement-related mesoscopic activity emerges across M1 and cerebellum during skill learning. ***A***, *Left:* Schematic of recording electrode locations in M1 and contralateral cerebellum depicted from top; *Right:* Histological verification of recording location in cerebellum (three markers used: Iba1 (green, microglia), GFAP (pink, astrocytes), DAPI (blue, nuclei). Sagittal section shows cerebellar lobules and cortical layers. Scale bar: 1 mm. Electrode shank is marked by two arrows. ***B***, *Top:* Example time course and illustration of recording scheme in M1 and the cerebellum from a frontal-side view. *Bottom*: neural and forearm muscle activity for representative successful trials from days 1 and 8.***C***, *Left*: Spectrograms of example M1 and cerebellar channels. *Right*: Difference in 1–4 Hz cerebellum and M1 power from early training to late training. The gray lines represent the mean power from individual animals (*n* = 4 animals) and the black lines represent the mean ± s.e.m. *P*-values are from mixed-effects models. ***D***, Left: Coherograms from an example M1 and cerebellum LFP channel pair. Right: difference in 1–4 Hz M1-cerebellum coherence from early to late training sessions. The gray lines represent the mean coherence from individual animals (*n* = 3 animals) and the black lines represent the mean and s.e.m. *P*-values are from mixed-effects models. ***E***, 1–4 Hz filtered LFP from example M1 and cerebellum channels time-locked to reach events; individual trials with mean overlaid. Bar plots depict changes in ITC from early trials to late trials (*upper row* for M1 and *lower row* for cerebellum). The gray lines represent the mean ITC from individual animals (*n* = 4 animals) and the black lines represent the mean and s.e.m. *P*-values from mixed-effects models.

We also analyzed movement-related low-frequency LFP coherence between M1 and cerebellum LFPs and we found that this also increased with increased task proficiency (**Fig. 2D**; 0.18 ± 0.02 coherence on early days to 0.21 ± 0.01 on late days, mixed-effects model: *t*(5158) = 13.4, *P* = 4.5 × 10^−40^). These increases in LFP power and coherence were not solely a by-product of faster and more consistent movements, since these high LFP power and coherence were not present for fast trials early in training, which we checked in a subset of animals.

With training, reaching sub-movements also became precisely phase-locked to 1–4 Hz LFP signals in both M1 and cerebellum, consistent with what we would expect if this activity was involved in generating sub-movements within this task (**Fig. 2E**; significant increase in inter-trial coherence (ITC) of the M1 LFP locked to movement onset (*MO*): mixed-effects model: *t*(406) = 4.4, *P* = 1.5 × 10^−5^; pellet contact (*PC*, right at the time of grasp initiation): mixed-effects model: *t*(406) = 7.0, *P* = 9.2 × 10^−12^; and retract onset (*RO*): mixed-effects model: *t*(406) = 10.1, *P* = 1.5 × 10^−21^; cerebellum LFP locked to movement onset: mixed-effects model: *t*(650) = 4.5, *P* = 8 × 10^−6^, pellet touch: mixed-effects model: *t*(650) = 9.8, *P* = 1 × 10^−19^, retract onset: mixed-effects model: *t*(650) = 9.1, *P* = 9.1 × 10^−19^).

### Coordinated spiking activity emerges across M1 and cerebellum during skill learning

The emergence of coordinated low-frequency activity across M1 and cerebellum was also clearly observed in movement-related spiking activity across M1 and cerebellum. We quantified phase-locking of movement-related M1 and cerebellar spikes to 1–4 Hz LFP signals in each region by generating polar histograms of the LFP phase at which each spike occurred for a single unit and LFP channel (**Fig. 3A**). The non-uniformity of the distribution of phases (indicating phase-locking) was quantified using a Raleigh test of circular non-uniformity. We compared all task-related M1 and cerebellar units on day 1 and 5 to a representative LFP channel in M1 and cerebellum and observed an increase in the percentage of M1 and cerebellum units phase-locked to both M1 and cerebellum LFP signals with training (**Fig. 3B**; the black vertical dashed lines correspond to the *P* = 0.05 significance threshold of the natural log of the Z statistic; M1 unit – M1 LFP pairs: 39.1% day 1 to 59.9% day 5, *P* = 8 × 10^−6^, Kolmogorov–Smirnov test; M1 unit – cerebellum LFP pairs: 21.8–76.3%, *P* = 1.4 × 10^−30^, Kolmogorov–Smirnov test; cerebellum unit – M1 LFP pairs: 77.8– 88.1%, *P* = 0.3, Kolmogorov–Smirnov test; cerebellum unit – cerebellum LFP pairs: 40.9–86.0%, *P* = 2.3 × 10^−10^, Kolmogorov–Smirnov test). All the pairs showed a significantly increased phase-locking, except cerebellum unit–cerebellum LFP pairs, although they also trended in same direction over days. These results further suggest that coordinated low-frequency activity emerges across M1 and cerebellum during skill learning.

**Figure 3.**
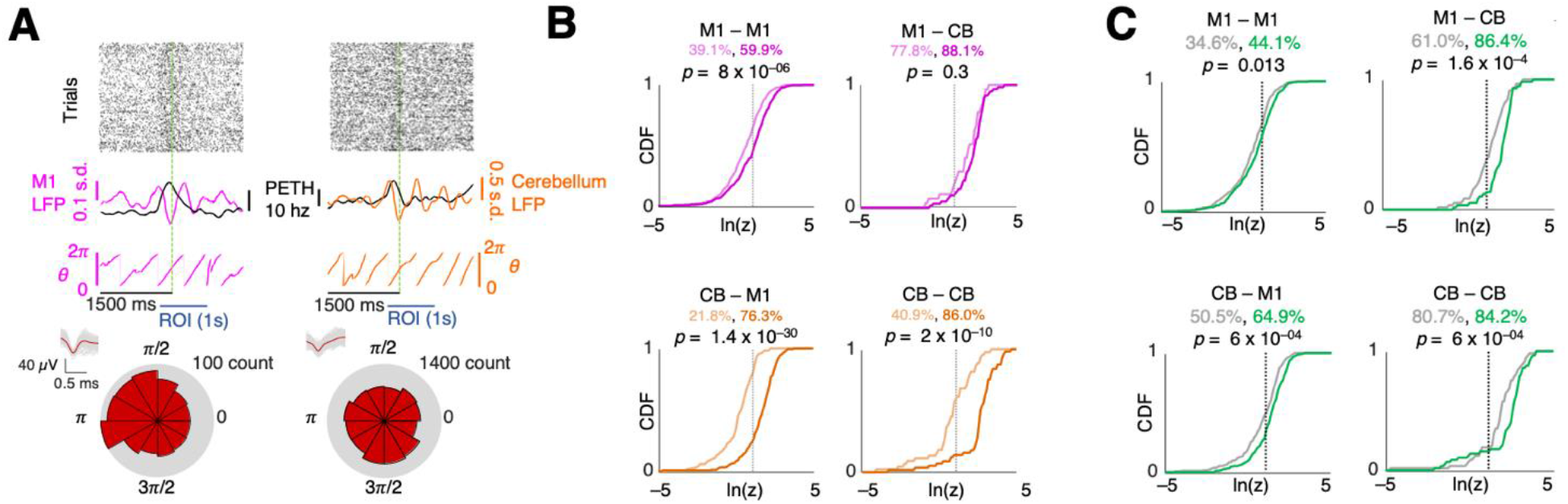
Coordinated spiking activity emerges across M1 and cerebellum during skill learning. ***A***, Illustration of spike locking to LFP phase for M1 unit-M1 LFP (left) and cerebellum unit-cerebellum LFP (right) pair examples. *Top*: raster plots of reach-centered spiking activity from example single units. *Middle*: 1–4 Hz filtered LFP overlayed with PETH from example unit. Below is the extracted phase from the filtered LFP. *Bottom*. Polar histograms of the spikes that occurred at various LFP phases. ***B***, Cumulative density functions (CDFs) of the z-statistic for every LFP-unit pair across and within each region. The vertical dotted lines indicate the significance threshold (*p* = 0.05). The percentage of the pairs with significant *p*-values is displayed. Lighter colors indicate early trials and darker is later. *n* = 280 M1 unit-LFP pairs on day 1, *n* = 298 M1 unit-LFP pairs on day 5, *n* = 46 cerebellum unit-LFP pairs on day 1, *n* = 73 cerebellum unit-LFP pairs on day 5. *P*-values derived using a Kolmogorov-Smirnov test. ***C***, Success/failure CDFs of the z-statistic for every LFP-unit pair within and across each region. The vertical dotted lines indicate the significance threshold (*p* = 0.05). The percentage of the pairs with significant *p*-values is displayed. Green indicate successful trials and gray is failures. *P*-values derived using a Kolmogorov-Smirnov test.

Next, we also explored these relations for successful and unsuccessful trials on day 5. We found that all four pairs showed significant M1 and cerebellar units’ phase-locking to 1–4 Hz M1 and cerebellum LFPs for successful trials (**Fig. 3C**; the black vertical dashed lines correspond to the *P* = 0.05 significance threshold of the natural log of the Z statistic; M1 unit – M1 LFP pairs: 34.6% for unsuccessful trials versus 44.1% for successful trials, *P* = 0.013, Kolmogorov–Smirnov test; M1 unit – cerebellum LFP pairs: 50.5–64.9%, *P* = 6 × 10^−4^, Kolmogorov–Smirnov test; cerebellum unit – M1 LFP pairs: 61.0–86.4%, *P* = 1.6 × 10^−4^, Kolmogorov–Smirnov test; cerebellum unit – cerebellum LFP pairs: 80.7–84.2%, *P* = 2.3 × 10^−10^, Kolmogorov–Smirnov test).

### Reorganization of neural dynamics in M1 and cerebellum with skill learning

We also investigated the consistency of single-trial population spiking activity by computing the correlations between single-trial neural activity and the trial-averaged template across all units in a session (**Fig. 4**). In early sessions, trial-to-trial neural firing was more inconsistent compared to later sessions, while later sessions were consistently associated with a stereotyped sequence of unit activations that also matched peri-event time histograms (PETH). This was observed in both M1 (**Fig. 4A**) and cerebellar (**Fig. 4C**) activity. Across the sessions from all rats, we observed a significant increase in template correlation among trials (**Fig. 4B, D**; linear mixed-effects model. M1: *t*(770) = 8.2, *P* = 5 × 10^−3^), cerebellum: *t*(648) = 5.4, *P* = 8 × 10^−8^) indicating that trial-to-trial variability in M1 and cerebellum neural activity reduced with skill consolidation.

**Figure 4.**
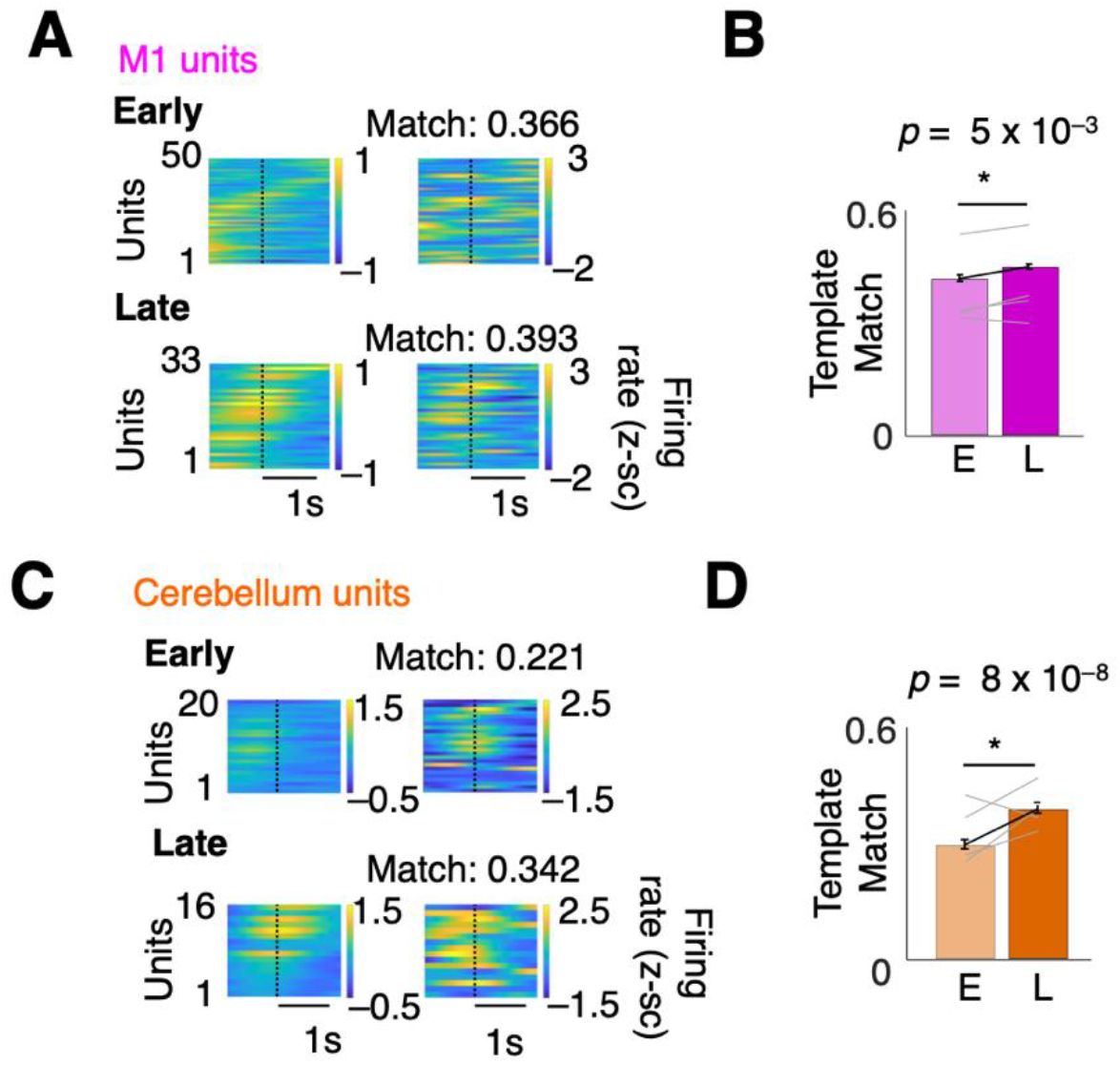
Changes in M1 and cerebellum neural dynamics with skill learning. ***A***, M1 successful trial averaged PETH from an example rat (*left*) and single trial PETH example (*right*) for early (*top*) and late (*bottom*) training sessions. ***B***, M1 PETH template match over training. Bars indicate mean ± s.e.m. over trials. Gray lines indicate average per animal (*n* = 4 animals). *P*-values are from mixed-effects models. ***C***, Cerebellum successful trial-averaged PETH from an example rat (left) and single trial PETH example (*right*) for early (*top*) and late (*bottom*) training sessions. ***D***, Cerebellum PETH template match over training. Bars indicate mean ± s.e.m. over trials. Gray lines indicate average per animal (*n* = 4 animals).

### Skilled movement representation in M1 and cerebellum

Lastly, we explored the representation of successful and failed reaches in M1 and cerebellum. We used Gaussian-process factor analysis (GPFA) to find low-dimensional neural trajectory representations of population spiking activity in M1 and cerebellum on individual trials^3,51^ (**Fig. 5A**) and then compared trajectories for successful and unsuccessful trials in early and late learning. We observed a difference between trajectories for successful and unsuccessful trials in M1 and cerebellum. To compare successful and unsuccessful trials we computed the correlation between the mean neural trajectory for successful trials, that is, the ‘successful template’, and each individual trial’s neural trajectory (**Fig. 5B**) during the period from 250 ms before movement onset until 250ms after retract onset (**Fig. 5C**). This period encompassed the movement onset and pellet contact for grasping and retraction of the forelimb. Since trials differed in the duration of this period, we interpolated trajectories such that they were all the same length. Neural trajectories for unsuccessful trials had significantly lower correlation than successful trials (**Fig. 5C**, Early M1: *P* = 8.5 × 10^−12^, Early cerebellum: *P* = 0.02, Late M1: *P* = 1.1 × 10^−8^, Late cerebellum: 6.6 × 10^−3^ mixed-effects model with Bonferroni correction for multiple comparisons). Together with Figure 3c, this provided further evidence that accurate reach-to-grasp task execution has M1 and cerebellar reliance.

**Figure 5.**
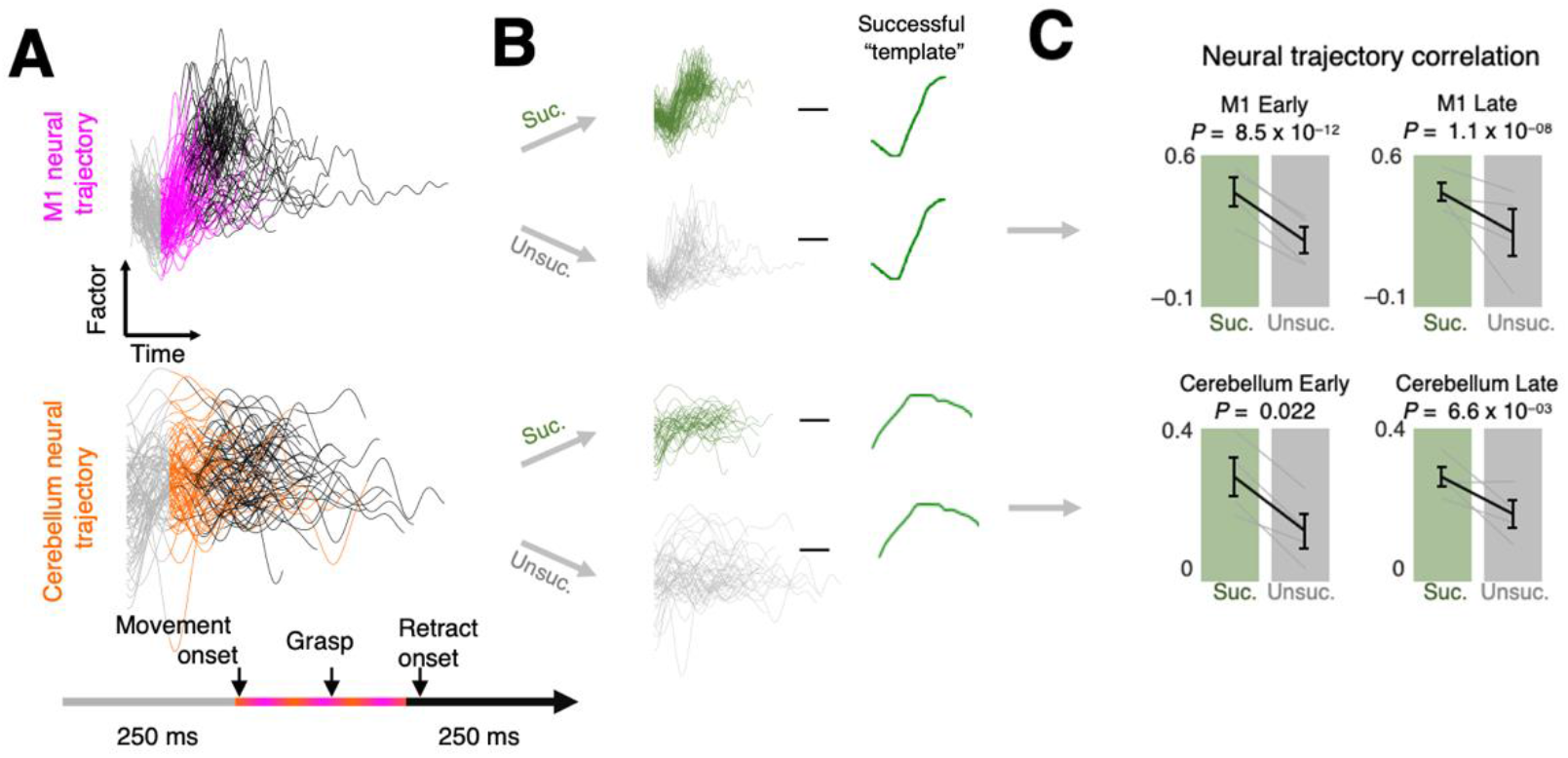
Skilled movement representation in M1 and cerebellum. ***A***, Example GPFA neural trajectories from late training sessions for M1 (*top*) and cerebellum (*bottom*) in a single animal. ***B***, Illustration of the process of comparing factor trajectories from successful and unsuccessful trials to the template (mean successful trajectory). ***C***, Deviation from the template for M1 (*top*) and cerebellum (*bottom*) factors. Gray lines represent individual animals (*n* = 4 animals), and the black line is mean ± s.e.m. across animals. *P*-values are from mixed-effects models.

## Discussion

In summary, we found that coordinated low-frequency activity emerged across M1 and cerebellum, which was linked to the emergence of faster reaching movements. We further found that coordinated spiking activity in both these regions was linked to accurate reach-to-grasp movements. Our work details the mesoscopic transmission in cortico-cerebellar networks and how it evolves with skill learning as well as how skilled reaching has a motor cortical and cerebellar cortical reliance.

### Role of M1 and cerebellum in motor skill learning and execution

M1 has a well-established role in motor learning as well as movement execution^52^. In particular, M1 is critical for generation of skilled dexterous movement^3,20,43,52^. Our work is consistent with this notion as we also see that M1 activity is different for successful reaches (**Fig. 3C, 4A** and **5**). M1’s projection to the cerebellum is thought to mediate fine-tuning of the movement. Cerebellar neurons in the cortex and in the deep nuclei are known to be modulated around several movement events. Perturbation of M1 input to cerebellum or direct manipulation of cerebellum itself is shown to delay movement initiation and to increase movement variability and duration^13,34,37,53^. Our work is also consistent with these observations as we found that precise, accurate movements had more consistent spiking activity in the cerebellum, and its coordination with LFOs in LFPs differed for successful and unsuccessful reaches (**Figs. 3C,4B** and **5**). Besides this role in increasing movement precision, cerebellar cortex is also theorized to contribute to task relevant dimensionality expansion that can aid in flexible computation and enhance learning^54–56^. This notion of dimensionality expansion was confirmed experimentally with the observations of high correlations among granule cells activity when mice expertly exerted pushing control over a manipulandum in a forelimb movement task^11^. This work also showed increase in emergent shared variance in M1 and cerebellar cells. Our increased M1-cerebellum LFP coherence with skill learning is similar to this observation. Neural network models of cortico-cerebellar networks show that cerebellar feedback improves rate of learning and cerebellar network also carries task representation^57^. Our experimental data supports this notion as well. We observed that M1-cerebellum LFP coherence increased with learning, and we observed movement-modulated units in the cerebellum. One of our observations also showed that M1 LFP-cerebellar units showed strong coordination in low-frequency range early on in training (**Fig. 3B**). This might suggest that cerebellar activity was critical during reach-to-grasp skill acquisition and is consistent with the notions of M1 being input-driven, and is also consistent with the cerebellar contributions to acquisition of skilled volitional movements^24,32^.

### Coordinated oscillatory dynamics across motor networks

One of our keys findings here is on low-frequency activity across M1 and cerebellum as an important marker of skilled motor control. We found evidence of such activity at the level of neural spiking and LFPs during the performance of dexterous task in rats. It is noteworthy that similar LFOs were recently shown to be disrupted in M1 post-stroke and tracked recovery^2^. This work also boosted M1 LFOs through electric stimulation to promote recovery. Recently, there has also been an interest in cerebellar stimulation for stroke recovery^58–60^, but a biomarker in cortico-cerebellar networks that can be target for closed-loop electric stimulation for stroke recovery is lacking. Future work can test if the LFOs we found in cortico-cerebellar networks of healthy animals with skill consolidation here can also serve as a biomarker for motor function during recovery from stroke. Mesoscopic biomarkers such as LFPs present a lower translational barrier in clinical populations.

Cortical LFOs can be used to decode reach-related activity and predict spiking phase across multiple behavioral states^9,61^. Such activity is also correlated with multiphasic muscle activations and timing of movements^5,8,9,62^. Recent work also suggest that oscillatory dynamics reflect an underlying dynamical system^5^. This previous work argues that LFOs represent an intrinsic property of motor circuits associated with precise movement control. Our findings extend this body of work by showing LFO dynamics in both M1 and cerebellum (**Fig. 2**). The exact origin of LFOs and underlying generators remains unknown. While reach-related LFOs may have involved striatum^3^ or thalamocortical activity^63^ so far, our results here raise the possibility of cerebellar involvement. Further work can probe interactions between M1 and the broader motor network to pinpoint the drivers of the electrophysiologic changes seen during skill learning.

## Materials and Methods

### Animal care and surgery

All procedures were in accordance with protocols approved by the Institutional Animal Care and Use Committee at the Cedars-Sinai Medical Center. Adult male Long Evans rats (n = 13, 250– 400 g; Charles River Laboratories) were housed in a 14-h/10-h light–dark cycle. All experiments were done during the light cycle. We used 8 rats for behavior only (**Fig. 1**) and 5 rats for behavior and physiology (**Figs. 2** to **5**; see **Supplementary Table 1** for details). No statistical methods were used to predetermine cohort size, but our sample sizes are similar to those reported in previous publications^3,50,64–66^ (3–9 animals per group). Animals were pair-housed prior to electrode implantation or behavioral training and then single-housed after to prevent damage to implants, or to implement food restriction, respectively.

All surgical procedures were performed using sterile techniques under 1%–4% isoflurane. Surgery involved cleaning and exposure of the skull and preparation of the skull surface using cyanoacrylate and then implantation of the skull screws for referencing and overall head-stage stability. The analgesic regimen included the administration of 0.1 mg per kg body weight buprenorphine, and 5 mg per kg body weight carprofen. Neural implanted rats were also administered 2 mg per kg body weight dexamethasone and 33 mg per kg body weight sulfatrim for 5 days. All neural implanted animals were allowed to recover for 5 days prior to further behavioral training. Ground and reference screws were implanted posterior to lambda contralateral to the recorded cerebellum, contralateral to the neural recordings. For M1 recordings, 32-channel arrays (33-μm polyamide-coated tungsten microwire arrays) were lowered to a depth of ∼1,200–1,500 μm in either the left or right M1 depending on handedness. These were implanted centered at 0.5 mm anterior and 3 mm lateral to the bregma^3,50^. For cerebellar recordings we used 32-64 channel tetrodes (Neuronexus, MI) or shuttle-mounted polytrodes (Cambridge Neurophysiology, UK). The probes were lowered into the cerebellar cortex through a craniotomy centered at 12.5 mm posterior and 2.5-3 mm lateral to bregma. Shuttle mounted probes were moved across days and recorded from depths of 1.5-4 mm. Our target regions were Simplex/ Crus I and Crus II areas of the cerebellum^67–69^. Activity in these areas has shown modulation during upper limb motor behaviors and in response to corticofugal fiber and forelimb stimulation. For the cerebellar recordings, we confirmed the location of electrode tips either through: (i) Staining with the orange/red fluorescence stain DiI (ThermoFisher Scientific) or (ii) three markers of Iba1 (microglia), GFAP (astrocytes), DAPI (nuclei) as shown in figure 2a (also see details below).

## Supplementary Table 1

**Supp. Table 1.**
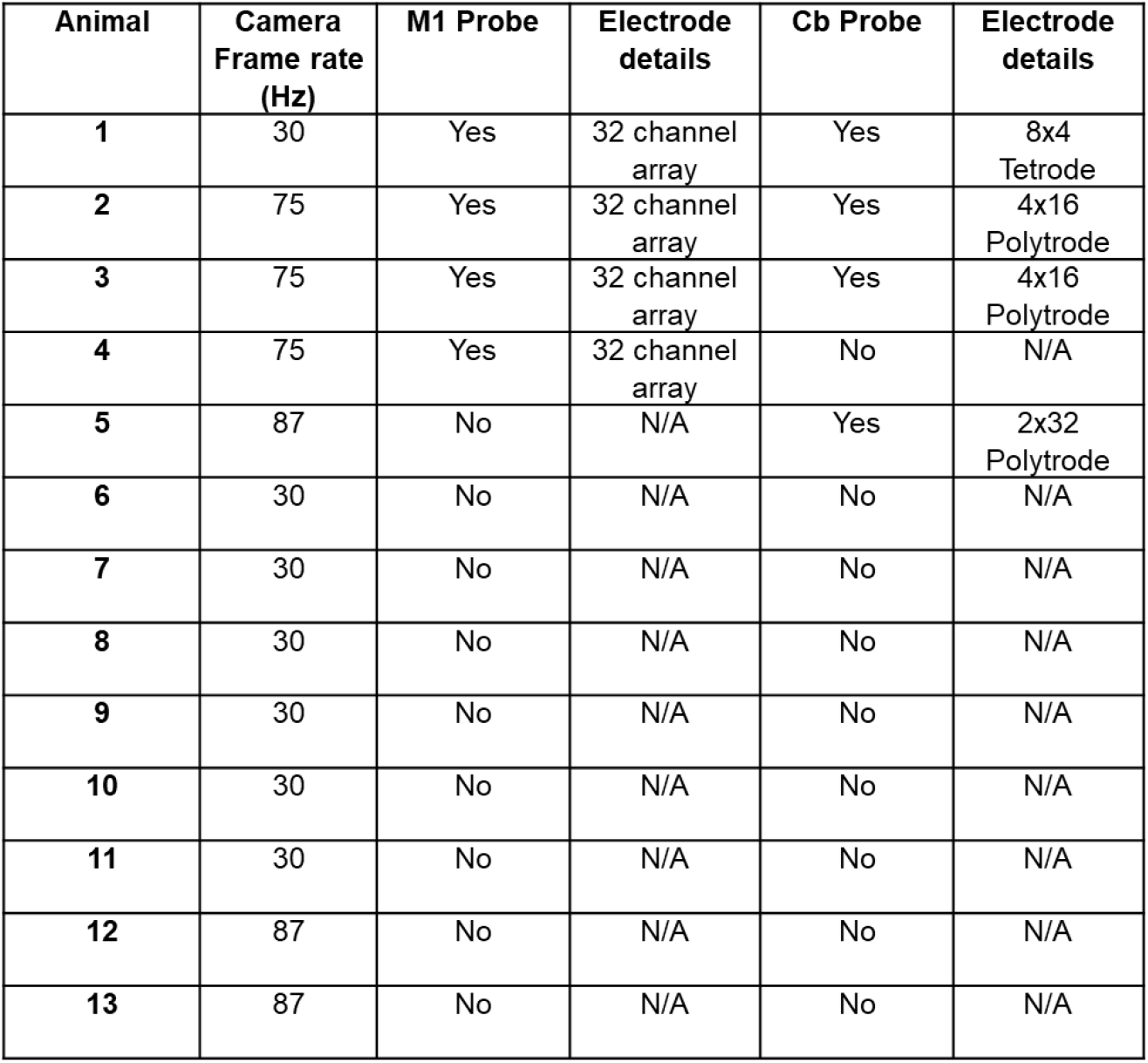
Number of rats used for experiments. Tabulated list of animals and behavioral monitoring camera frame rates and electrode used (see columns).

### Behavior

#### Training

Rats were acclimated to the behavioral box for at least 2 days and then exposed to a reach-to-grasp task for 5-10 trials to establish hand-preference before neural probe implantation. Probe implantation was performed in the contralateral M1 and ipsilateral cerebellum to the preferred hand. Rats were allowed to recover for at least 5 days before the start of experimental sessions. During behavioral assessments, we monitored the animals and ensured that their body weights did not drop below 90% of their initial weight. We used an automated reach-box, controlled by custom MATLAB scripts and an Arduino microcontroller. This setup requires minimal user intervention, as described previously^41^. Each trial consisted of a pellet dispensed on the pellet tray, followed by an alerting beep indicating that the trial was beginning. They then had 15 s to reach their arms through the slot, grasp and retrieve the pellet. A real-time ‘pellet detector’ using an infrared detector centered over the pellet was used to determine when the pellet was moved, which indicated that the trial was over, and the door was closed. All trials were captured by video through a camera placed on the side of the behavioral box. The camera was synced with the electrophysiology data either using Arduino digital output or directly through TTL pulses to the TDT RZ2 system. In electrode implanted animals the video frame rate ranged from 75-87 Hz aside from 1 animal for which the framerate was 30 Hz (see **Supp. Table 1**). For behavior-only animals the framerate was 30Hz aside from two animals for which the framerate was 87 Hz.

#### Behavioral Testing

Rats began behavioral testing training 5 days after surgery by performing the same reach-to-grasp task. Electrophysiology recordings were taken throughout the full extent of the testing which consisted of one to two sessions of 60-100 trials per day for 5 days. Typically, each day would consist of a session of 100 trials followed by a session of 60 trials. Sessions within a day were spaced by a 2-hour resting block.

#### Behavioral analysis

Behavioral analysis was done based on video recorded during experimental sessions. Reach videos were viewed and manually scored to obtain trial success, hand position and time points for reach onset, pellet contact and retract onset. To characterize motor performance, we quantified pellet retrieval success rate (percentage of pellets successfully retrieved into the box) and reach duration (time from reach onset to retract onset).

### In vivo electrophysiology

Units and LFP activity were recorded using a 128-channel TDT-RZ2 system (Tucker-Davis Technologies). Spike data were sampled at 24,414 Hz and LFP data at 1,017.3 Hz. ZIF (zero insertion force) clip-based digital head stages from TDT were used that interface the ZIF connector and the Intan RHD2000 chip that uses 192x gain. Behavior-related timestamps (that is, trial onset, trial completion) and video timestamps (that is, frame times) were sent to the RZ2 analog input channel using an Arduino digital board and synchronized to the neural data.

### Neural data analysis

Analyses were conducted using custom-written scripts and functions in MATLAB 2018a (MathWorks, MA).

#### Local field potential (LFP) analyses

Artifact rejection was first performed on LFP signals to remove broken channels and noisy trials. LFPs were then z-scored and median referenced separately for M1 and cerebellum. There was excessive noise detected in all cerebellum channels in 1 of the 4 animals with simultaneous M1 and cerebellum recordings. Hence, the cerebellar LFP activity from that animal was excluded from analysis. A fifth animal with only cerebellum implants was included in the cohort. LFP power was calculated on a trial-by-trial basis and then averaged across channels and animals, with wavelet decomposition using the EEGLAB^70^ function ‘newtimef’. To characterize coordination of activity across regions, we measured changes in movement-related spectral coherence between LFP channels in M1 and cerebellum. For learning comparisons, coherence was measured for the same channels on early and late days, and specifically for channels with an increase in power of 0.5 baseline-normalized unit from early to late days. Strong coherence in a specific frequency band indicates a constant phase relationship in that frequency between two signals and is theorized to indicate increased communication between regions^3,71,72^. M1-cerebellum LFP coherence was calculated for each pair of channels using the EEGLAB function ‘newcrossf’ with 0.1s windows moving by 0.01s.

To determine whether the emergence of coordinated low-frequency activity during training was attributable solely to faster movements, we compared LFP power and coherence between ‘fast’ trials (trials with a movement duration less than 300 ms) on day 1 and 2 versus day 4 and 5.

In several instances, we filtered the LFP signals to isolate and display the low-frequency (1–4 Hz) component of the signal (**Figs. 2** and **3**). Filtering was performed using the EEGLAB function ‘eegfilt’^70^. In addition to display purposes, we also used filtered LFP to characterize the phase-locking of spiking activity specifically to low-frequency LFP signals. For this we used the Hilbert transform linear operator (MATLAB) to extract the phase information from low-frequency filtered LFP signals (**Fig. 3**).

To quantify phase-locking of LFP signals to specific sub-movements (movement onset, pellet contact and retract onset), we calculated the ITC of LFP signals across trials time-locked to these sub-movements (**Fig. 2d**). ITC was measured and compared for the same channels on early and late days across all channels (except those removed due to noise). ITC was computed using the EEGLAB function ‘newtimef’^70^.

#### Spiking analyses

Spike data was sorted offline after local median subtraction. The threshold for spiking activity was set offline using the automated spike detection of Spyking Circus, and waveforms and timestamps were stored for any event that crossed that threshold. Sorting was performed using a principal component analysis (PCA)-based method followed by manual inspection and sorting. Clearly identified units were selected for this analysis. All units were analyzed and not sorted into cell type based on waveform shape. However, activity that was unambiguously multi-unit was removed. Behavior-related timestamps (trial onset and trial completion) were sent to the RZ2 analog input channel using an Arduino digital board and synchronized to neural data.

#### Unit modulation and spike-LFP phase analysis

Spikes were binned at 25 ms and time locked to behavioral markers. For visualization purposes, the peri-event time histogram (PETH) was estimated by the MATLAB ‘fit’ function using smoothing spines. To determine if a unit was significantly modulated during movement, a baseline firing rate mean and standard deviation was taken within the period –4s to –2s from reach onset. If the mean firing rate in the period from –350ms to –850ms relative to reach onset differed from the baseline mean by more than 1.25 baseline standard deviations the unit was categorized as a reach-modulated unit.

To characterize low-frequency spiking activity, we generated histograms of the LFP phases at which each spike occurred for a single unit to a single LFP channel filtered in the 1–4 Hz band in a 1-s window around movement (–250 ms before to 750 ms after movement onset) across all trials of a session (**Fig. 3**). For learning comparisons, all units were compared to the same selected M1 and DLS LFP channel on day 1 and 5. These histograms were generated for each unit–LFP channel pair both within and across regions. For every pair, we then calculated the Rayleigh’s z-statistic for circular non-uniformity. These z-statistics were then used to calculate the percentage of significantly non-uniform distributions across unit–LFP pairs with a significance threshold of P = 0.05 (**Fig. 3**). A significantly nonuniform distribution signifies phase preference for spikes of a unit to an LFP signal. This process was also performed to compare the successful and unsuccessful trials of day 5 (**Fig. 3C**).

#### Single trial to template correlation

Spikes from –4s to 4 s around pellet touch were binned at 20 ms, smoothed with a Gaussian kernel with a standard deviation of 60 ms and then z-scored. Binned, smoothed and standardized spike counts within the period of –1s to 1.25s for all units of a single trial were then concatenated into one long vector. The correlation (measured using Pearson’s r) between each concatenated single trial neural activity and the mean template (mean of all successful trials) was computed and the mean correlation for each session was reported (**Fig. 4**).

#### GPFA neural trajectory analyses

To characterize single-trial representations of population spiking activity we used GPFA^3,51^ to find low-dimensional neural trajectories, which consisted of the first two factors, for each trial. GPFA analyses were carried out using the MATLAB based graphical user interface DataHigh (version 1.2)^73^, 10 ms time bins and a dimensionality of 5. We determined the magnitude of deviation for each individual trial trajectory from the mean trajectory across all successful trials by taking the absolute value of the difference between the trajectory of each trial and the mean trajectory across all trials (**Fig. 5B, C**; computed in each dimension independently). This was performed specifically for the period between 250 ms before movement onset and until 250 ms after retract onset. Since this duration varied across trials, we interpolated each trial such that every epoch (reach onset to touch and touch to retract onset) of each trial was the same length and then calculated the average deviation.

### Immunohistochemistry

After all experiments, rats were anesthetized and transcardially perfused with 1% phosphate-buffered saline, followed by phosphate-buffered 4% formaldehyde (PFA). The harvested brains were post-fixed for 72 h in PFA and immersed in 30% sucrose. For immunofluorescence staining (**Fig. 2A**), sagittal cerebellar tissue cryostat sections (40 μm) were washed 3x in 1x Tris Buffered Saline (TBS), followed by antigen retrieval with 0.1N hydrochloric acid (HCl). After 3 more washes in 1xTBS, sections were blocked with 5% Normal Donkey Serum (NDS) in 0.1% TBS-T(Triton) for 1 hour. Sections were then incubated in primary antibodies for astrocytes and microglia overnight. The next day sections were washed 3 times in 1xTBS and then incubated with fluorescent secondary antibodies for 2 hours. Sections were then washed 3 times in 1xTBS and incubated with 300nM DAPI in 1xTBS for 7 min, before coverslipping with mounting media (ProLongTM Glass Antifade Mountant, ThermoFisher cat# P36980). Primary antibodies used are 1:1000 Rat-anti-GFAP(ThermoFisher cat #13-0300) and 1:1000 Rabbit-anti-IBA1(Wako cat #019-19741). Secondary antibodies used are 1:250 Alexa FluorTM 647 Donkey-anti Rat (Jackson cat# 712-605-153) and 1:1000 Alexa FluorTM 488 Donkey-anti Rabbit 488(ThermoFisher cat # A-21206). Fluorescent sections were imaged with a BZ-X700 Keyence microscope.

### Statistical analysis

The linear mixed-effects model (implemented using MATLAB ‘fitlme’) was used in this study. Using these models accounts for the fact that units, channels or trials from the same animal are more correlated than those from different animals; thus, it is more stringent than computing statistical significance over all units, channels or trials^3,74^. We fitted random intercepts for each rat and reported the p values for the regression coefficients associated with successful or unsuccessful outcome, early (that constituted days 1 and 2) or late (that constituted days 4 and 5) learning, or training session. Linear mixed effects models was used for testing significance in **Figs 1D,E**; **2C-E**; **4B,D**; and **5C**. Two-sample Kolmogorov–Smirnov tests were used to test whether spike-LFP phase-locking values on days 1 and 5 came from the same distribution (**Fig. 3C**). All statistical analyses were implemented within MATLAB. We fitted random intercepts for each rat and reported the p values for the regression coefficients associated with successful or unsuccessful outcome, early or late learning, or training session.

## Acknowledgements

We thank Celine Riera and Joshua Burda for resources and reagents for histology. We thank Keshav Suresh for assistance with immunohistochemistry. This research is supported by American Heart Association (postdoctoral fellowship 897265 to A.A. and career development award 847486 to T.G.), National Institutes of Health (R00NS097620 to T.G.), National Science Foundation (CAREER award 2048231 to T.G.) and Cedars-Sinai Medical Center. A.A. also received support from a Cedars-Sinai’s Center for Neural Science and Medicine postdoctoral fellowship.

## Author Contributions

P.J.F. and T.G. designed the study. P.J.F., A.A., A.W.F. and N.P.D. carried out the electrophysiology experiments. A.A., A.W.F., N.P.D. and T.G. performed the surgical procedures. P.J.F. carried out the analysis. A.A., A.W.F., N.P.D., R.S. and P.R.R. provided resources and assisted with analysis. A.W.F., N.P.D., R.S. and P.R.R. carried out histology. P.J.F. and T.G. wrote the manuscript. A.A. and A.W.F. edited the manuscript.

