## Supplemental Table 1 for "Emergent low-frequency activity in cortico-cerebellar networks with motor skill learning"

### Supplementary Table 1

| Animal | Camera Frame rate (Hz) | M1 Probe | Electrode details | Cb Probe | Electrode details |
|---|---|---|---|---|---|
| **1** | 30 | Yes | 32 channel array | Yes | 8x4 Tetrode |
| **2** | 75 | Yes | 32 channel array | Yes | 4x16 Polytrode |
| **3** | 75 | Yes | 32 channel array | Yes | 4x16 Polytrode |
| **4** | 75 | Yes | 32 channel array | No | N/A |
| **5** | 87 | No | N/A | Yes | 2x32 Polytrode |
| **6** | 30 | No | N/A | No | N/A |
| **7** | 30 | No | N/A | No | N/A |
| **8** | 30 | No | N/A | No | N/A |
| **9** | 30 | No | N/A | No | N/A |
| **10** | 30 | No | N/A | No | N/A |
| **11** | 30 | No | N/A | No | N/A |
| **12** | 87 | No | N/A | No | N/A |
| **13** | 87 | No | N/A | No | N/A |

**Supp. Table 1. Number of rats used for experiments.** Tabulated list of animals and behavioral monitoring camera frame rates and electrode used (see columns).
